# Quantitative modeling of EGF receptor ligand discrimination via internalization proofreading

**DOI:** 10.1101/2023.05.09.539827

**Authors:** Jaleesa A Leblanc, Michael G Sugiyama, Costin N Antonescu, Aidan I Brown

## Abstract

The epidermal growth factor receptor (EGFR) is a central regulator of cell physiology that is stimulated by multiple distinct ligands. Although ligands bind to EGFR while the receptor is exposed on the plasma membrane, EGFR incorporation into endosomes following receptor internalization is an important aspect of EGFR signaling, with EGFR internalization behavior dependent upon the type of ligand bound. We develop quantitative modeling, both kinetic and with spatial details, for EGFR recruitment to and internalization from clathrin domains and competition with ligand unbinding from EGFR. We find that a combination of spatial and kinetic proofreading leads to enhanced EGFR internalization ratios in comparison to unbinding differences between ligand types. Various stages of the EGFR internalization process, including recruitment to and internalization from clathrin domains, modulate the internalization differences between receptors bound to different ligands. Our results indicate that following ligand binding, EGFR may encounter multiple clathrin domains before successful recruitment and internalization. The quantitative modeling we have developed describes competition between EGFR internalization and ligand unbinding and the resulting proofreading.

## 1. Introduction

Receptor proteins on the plasma membrane enable signals, carried by extracellular molecules (ligands) that bind to the receptor, to be transduced to the cell interior [1]. The epidermal growth factor receptor (EGFR) is a receptor tyrosine kinase that mediates activation of a range of signaling pathways that regulate many aspects of cell physiology [2, 3, 4]. EGFR also plays a key role for tumor progression, representing a drug target and a drug resistance challenge for many cancers [5, 6, 7].

Ligands bind to EGFR on the plasma membrane, with recent work suggesting that EGFR ligand binding is favored for receptors localized to tetraspanin nanodomains [8]. While unliganded EGFR can form dimers, ligand-bound EGFR form distinct dimers and oligomers, which play an important role in EGFR signaling [9, 10]. Phosphorylation of intracellular EGFR domains, while EGFR is on the plasma membrane, is a key step for signal transduction [11, 12, 13].

Ligand-bound EGFR can be recruited to a clathrin domain or stabilize a nascent clathrin domain [8, 14, 15, 16, 17]. EGFR can be endocytosed upon generation of membrane curvature to form clathrin-coated pits, a process which can lead to formation of intracellular vesicles [18, 19]. While EGFR may be internalized via other pathways, under a range of conditions, clathrin-mediated endocytosis is the primary internalization pathway for EGF receptors at lower physiological EGF concentrations [18, 19]. EGFR signaling occurs both from EGFR on the plasma membrane and in endosomes following endocytosis [19, 20, 21, 22, 23, 24]. From endosomes, receptors are either transported to lysosomes for degradation or recycled back to the plasma membrane [25].

EGFR internalization from the plasma membrane into endosomes leads to EGFR signaling modes that are distinct from EGFR signaling from the plasma membrane [18, 19, 20, 22, 23, 24]. As signaling activity of EGFR in endosomes is lost when the ligand dissociates [23], ligand binding is important for endosome-based EGFR signaling.

EGFR is stimulated by at least seven distinct ligands: high-affinity ligands epidermal growth factor (EGF), heparin-binding EGF-like growth factor (HB-EGF), transforming growth factor-*α* (TGF-*α*), and betacellulin (BTC), which bind the cell surface with affinities of apparent dissociation constant *K*_d_ = 0.1 – 1 nM; and low-affinity ligands epiregulin (EREG), epigen (EPGN), and amphiregulin (AREG), which bind 10- to 100-fold more weakly [26, 27]. Although all bind to EGFR, these different ligands stimulate distinct responses, differentially activating various signaling pathways [28, 29, 30, 31]. There are many examples of specific cellular responses to different ligands [30], such as EGF leading to an epithelial to mesenchymal transition while AREG causes proliferation of colon cancer cells [32]; and AREG and TGF-*α* stimulation both inducing kidney cell mitosis, but only AREG impacting E-cadherin distribution [33]. While the ERK/MAPK signaling pathway is readily activated at low stimulating ligand concentration by EGF, HB-EGF, TGF-*α*, and BTC, the Akt, PLCg and STAT3 signaling pathways respond differently to EGF, HB-EGF, TGF-*α*, and BTC [29].

There has been substantial effort to understand the mechanistic underpinnings of how different ligands lead to different signaling pathway activation and cellular responses. For example, different ligands lead to distinct EGFR dimers with varying interaction strength and lifetime [26] and different degrees of EGFR trafficking to the nucleus [34]. We will focus on ligand-specific differences in receptor internalization into endosomes. HB-EGF and BTC lead to high internalization, EGF and TGF-*α* to somewhat lower internalization, and EPGN and AREG to the lowest internalization [25]. Internalized EGFR can be recycled to the plasma membrane or lysosomally degraded, with different ligands more likely to induce EGFR degradation: EGF, HB-EGF, and BTC cause substantial degradation and TGF-*α*, EREG, and AREG largely induce recycling to the plasma membrane [25, 29]. Degradation and intracellular retention of EGFR align with signaling persistence: TGF-*α*, EPGN, and AREG cause persistent signaling as the receptors they bind to are typically recycled to the plasma membrane, while EGF, HB-EGF, and BTC cause more transient signaling as they lead to EGFR degradation, lowering plasma membrane EGFR levels until replenished by protein synthesis [25]. The likelihood of receptor degradation with different ligands also largely aligns with internalization differences [19], ligand dissociation rates at endosomal pH (slower dissociation leads to more degradation) [25, 35], and ubiquitination and phosphorylation persistence on EGFR in endosomes (persistent modification leads to more degradation) [25].

As EGFR internalization into endosomes facilitates important signaling modes distinct from EGFR signaling modes from the plasma membrane and different EGFR ligands induce varying degrees of EGFR internalization, we explore a possible mechanism behind the variation in EGFR internalization due to binding of different ligands.

‘Kinetic proofreading’ describes a class of mechanisms that use nonequilibrium conditions to reduce the error rates or enhance discrimination between stimuli for various cellular and biochemical processes. While initially envisioned to increase accuracy of ‘reading’ or copying processes such as DNA replication or protein synthesis [36, 37], kinetic proofreading principles are understood to apply to a broader class of phenomena, particularly signaling and recognition processes [38, 39, 40, 41] such as those of receptors on the plasma membrane. To improve copying or signaling fidelity, these kinetic proofreading processes utilize multiple stages and repeated cycles to enhance the difference between ‘right’ and ‘wrong’ substrates. With multiple stages each discriminating between the substrate types, the combined effect is to enhance the overall difference in outcome between substrate types beyond the performance of an equilibrium process.

Recently, a distinct spatial proofreading mechanism was proposed [42], with the diffusive search time between a binding event at one location and a reaction event at another location allowing discrimination between ligands with different unbinding rates. For sufficiently distinct unbinding rates and an appropriate timescale for diffusive search, a receptor stimulated by a slower-unbinding ligand can be far more likely than a receptor stimulated by a faster-unbinding ligand to reach the final reaction location before ligand unbinding.

We propose that the EGFR internalization process following ligand binding, including distinct signaling modes due to interaction with clathrin-coated pits and endosomes, applies the principles of kinetic and spatial proofreading to improve ligand discrimination. We develop two related quantitative models for EGFR dynamics, one solely kinetic and another involving receptor diffusion on the plasma membrane. We find that the internalization ratio for EGFR stimulated by different EGFR ligands exceeds the unbinding rate ratio between the ligands for physiological parameters. This increased differentiation between EGFR ligands enables greater ligand specificity in EGFR signaling activity.

## 2. Results

### 2.1. Kinetic model

Our EGF receptor model considers receptors immediately following ligand binding, ending when the receptor is internalized into the cell during endocytosis of a clathrin domain or the ligand unbinds from the receptor. A kinetic version of this model is shown in Fig. 1A, with *k*_c_ the rate of a ligand-bound receptor *R*_L_ entering a clathrin domain, *k*_-c_ the rate of a clathrin-localized receptor *R*_C_ exiting a clathrin domain, *k*_i_ the rate of a clathrin-localized receptor *R*_C_ internalizing via endocytosis, and *k*_u_ the rate of a ligand unbinding from a receptor (inside *R*_C_ or outside *R*_L_ a clathrin domain).

**Figure 1.**
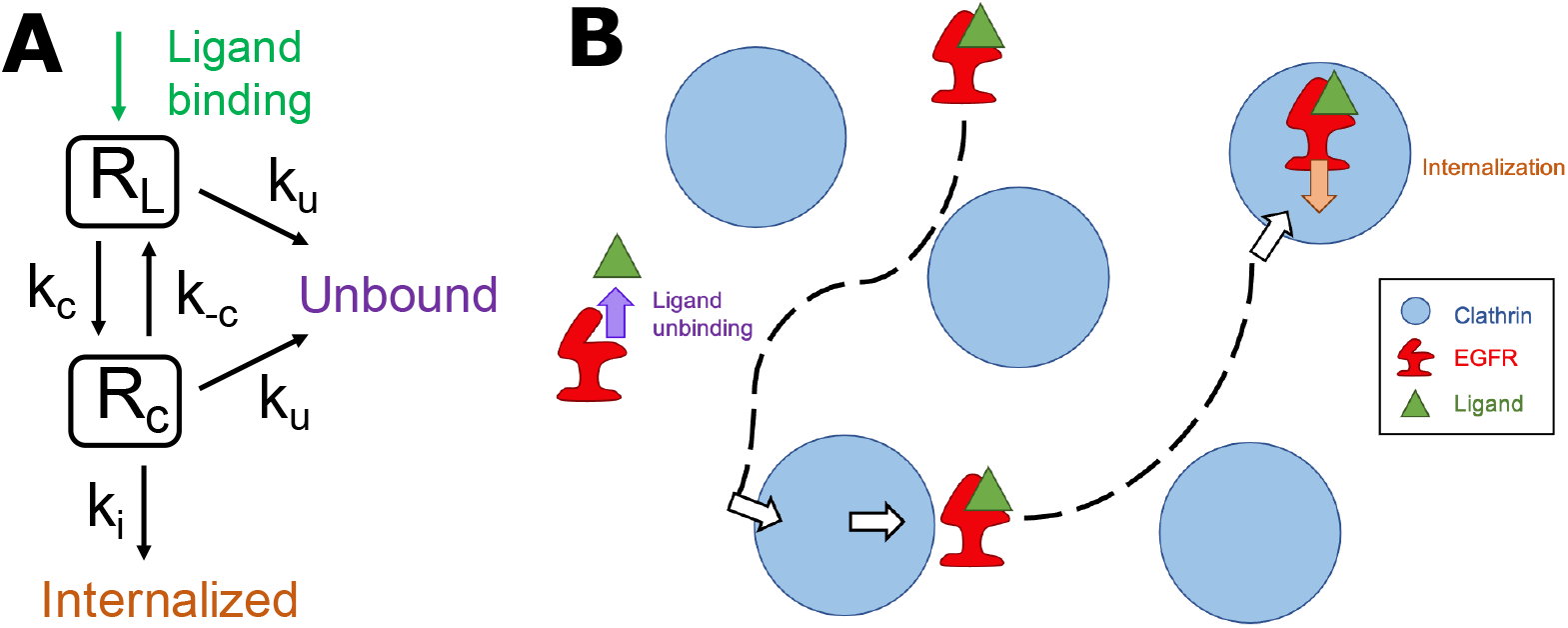
(A) Kinetic model of EGFR. Receptors begin following ligand binding in the ligand-bound state outside of clathrin domains (*R*_L_). These receptors can enter a clathrin domain (state *R*_C_) with rate *k*_c_ or have the ligand unbind at rate *k*_u_. Receptors in a clathrin domain can leave the domain with rate *k*_-c_, internalize at rate *k*_i_, or have the ligand unbind at rate *k*_u_. Ligand unbinding or receptor internalization are terminal states. (B) Schematic of spatial model of EGFR. Receptors (red) begin following ligand (green) binding outside of clathrin domains (blue). Receptors diffuse to encounter clathrin domains and within clathrin domains, and cross an energy barrier to enter or exit clathrin domains. Ligand unbinding (occurring both within and outside of clathrin domains) and internalization from within clathrin domains terminate the receptor trajectory.

For a receptor beginning in state *R*_L_, the probability *P*_i_ that a receptor is internalized is (see Appendix for derivation)

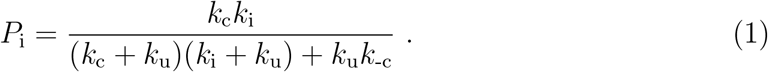

We estimate the clathrin entry rate *k*_c_ ≈ 0.1 s^−1^, which is substantially slower than the estimated diffusion-limited entry rate [43] of ≈ 1 s^−1^ (for EGFR with diffusivity 0.2 *μ*m^2^*/*s [8, 9, 44, 45, 46, 47] to clathrin domains of 50 nm radius representing 1% of the plasma membrane surface [48]). *k*_-c_ ≈ 0.05 s^−1^ and *k*_i_ ≈ 0.05 s^−1^ are estimated from clathrin domain dynamics [48]. See the Appendix for the details of these parameter estimates from experimental observations.

### 2.2. Proofreading

In Fig. 2A the internalization probability *P*_i_ is shown as the ligand unbinding rate *k*_u_, which will be distinct for different ligands, is varied. Figure 2A shows *P*_i_ for the estimated EGFR parameters as well as with each parameter *k*_c_, *k*_-c_, or *k*_i_ increased or decreased relative to the estimated parameters by an order of magnitude. At low *k*_u_ the internalization probability is flat at *P*_i_ ≈ 1, as the unbinding rate is too small to compete with internalization. As the unbinding rate increases, the internalization probability decreases, eventually reaching a limiting power law for high *k*_u_ of 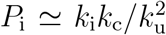. This power law, reached only for sufficiently high unbinding rates, corresponds to receptors that effectively make a single attempt to enter a clathrin domain and then internalize or have the ligand unbind, without sufficient time for another internalization opportunity. EGFR internalization switches from a nearly certain event for low *k*_u_ to a rare event for *k*_u_ that are two to three orders of magnitude larger.

**Figure 2.**
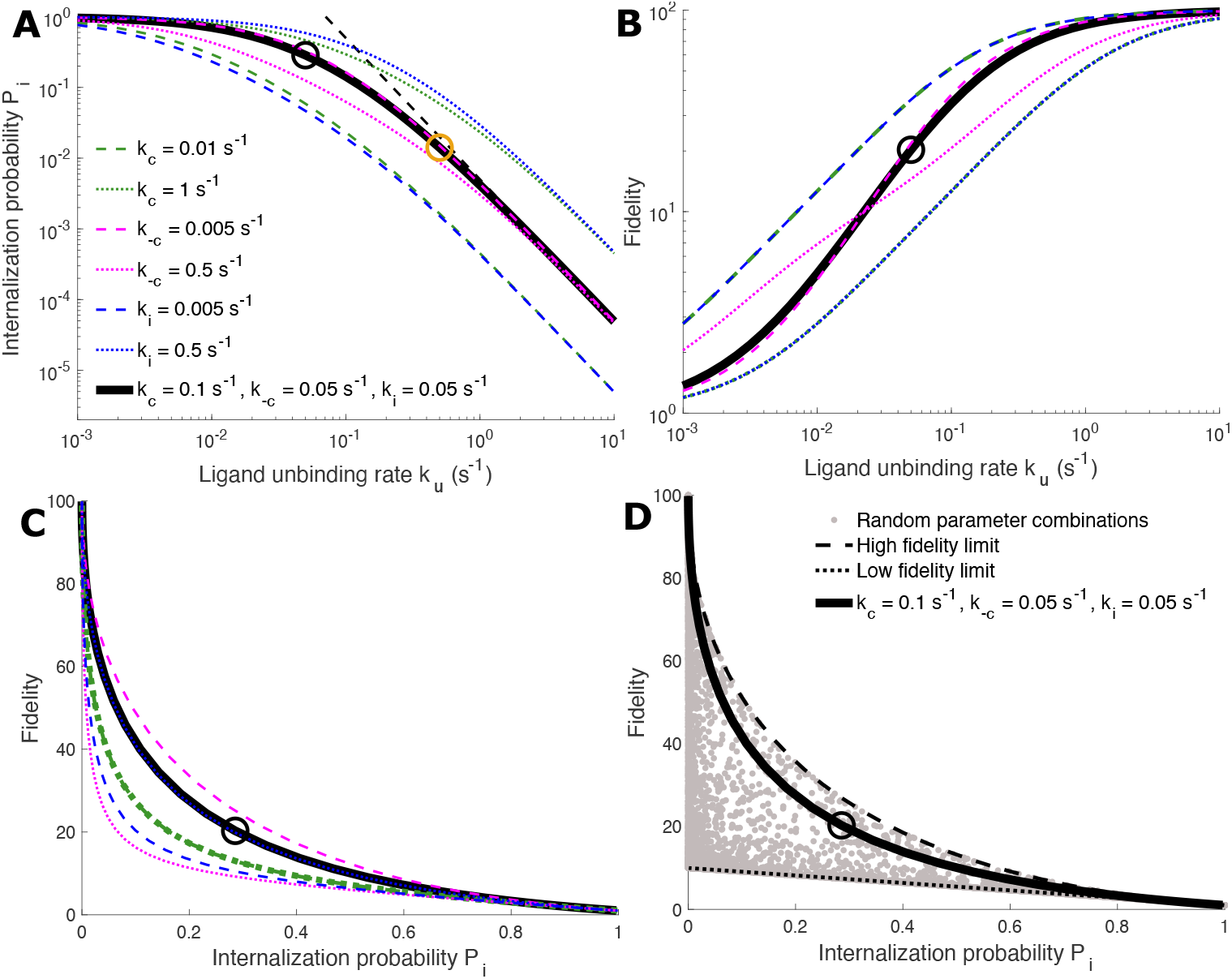
Internalization probability and fidelity with kinetic model. (A) Internalization probability *P*_i_ (Eq. 1) as ligand unbinding rate *k*_u_ is varied. Thick solid black curve shows *P*_i_ with estimated EGFR parameters (see text). Other curves have one parameter value changed compared to thick black curve, as indicated. Black circle shows *P*_i_ for *k*_u,EGF_ = 0.05 s^−1^ and orange circle shows *P*_i_ for *k*_u,EREG_ = 0.5 s^−1^. Black dashed line is the high *k*_u_ limiting power law 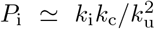. (B) EGFR internalization fidelity (Eq. 4) between a fast and slow unbinding ligand with ratio 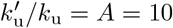 as the unbinding rate of the slow unbinding ligand *k*_u_ is varied. Black circle shows the fidelity for slow unbinding rate *k*_u,EGF_ = 0.05 s^−1^. (C) Fidelity vs internalization probability as *k*_u_ is parametrically varied. Curves in (B) and (C) take parameter values indicated in legend in (A). (D) Limits of fidelity vs internalization probability relationship as *k*_u_ is parametrically varied. The high fidelity limit (black dashed curve) is Eq. 6 and the low fidelity limit (black dotted curve) is Eq. 5. Random parameter combinations (grey) is 10^4^ random, logarithmically-distributed, and independent samples for *k*_c_, *k*_i_, *k*_-c_, and *k*_u_ from the range [10^−4^, 10^3^] s^−1^.

The unbinding rate *k*_u_ varies between EGFR ligands. EGF has an unbinding rate experimentally measured as 0.02 – 0.04 s^−1^ [49], 0.04 s^−1^ [50], 0.055 s^−1^ [51], 0.06 s^−1^ [52], 0.062 s^−1^ [53], and 0.066 s^−1^ [54]. Selecting an intermediate value of EGF unbinding from EGF receptors of *k*_u,EGF_ = 0.05 s^−1^, the internalization probability of EGFR following EGF binding is approximately 29% (black circle in Fig. 2A).

EGFR binding affinity varies for other ligands. Epiregulin (EREG) has a weak EGFR affinity compared to other ligands [27], with at least 10× weaker affinity for EGFR compared to EGF [26, 27]. As EGFR ligands have the same binding motif [55], use similar binding modes [56], and ligand affinities can be accurately ranked by ligand-receptor interaction energies [56], we assume that binding rate constants are similar for different EGFR ligands and that unbinding rate differences are proportional to the affinity differences. Thus we take the EREG unbinding rate to be 10× faster than that of EGF, *k*_u,EREG_ = 0.5 s^−1^. This corresponds to a receptor internalization probability of approximately 1.4% (orange circle in Fig. 2A), which is approximately 20× less internalization than receptors bound to EGF.

Proofreading often describes the ratio between a ‘right’ and a ‘wrong’ product or signal. However, distinctions in EGFR response to stimulation by different ligands such as EGF and EREG do not clearly fit into categories of ‘right’ and ‘wrong’ as these EGFR ligands each send meaningful signals to a cell via EGFR. We will instead treat EGFR internalization as a signaling aspect with a propensity that differs between ligand types. We then calculate the fidelity [42], or ratio of receptor internalization probabilities for different ligand types, as a measure of the degree to which EGFR internalization can distinguish between stimulation by different ligand types. The EGFR internalization fidelity between a ligand with unbinding rate *k*_u_ and a ligand with unbinding rate 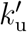 is then

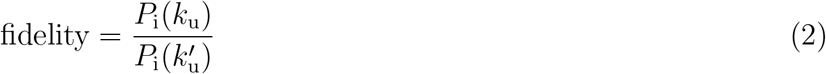

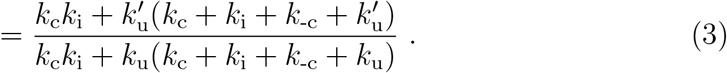

If the ratio between the unbinding rates is 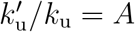, then

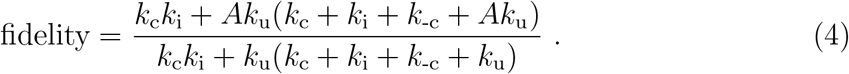

In Fig. 2B the fidelity is shown as the unbinding rate *k*_u_ is varied for unbinding rate ratio *A* = 10, corresponding to the estimated ratio between EGF and EREG ligands. For low *k*_u_, the fidelity approaches unity, as binding of both ligand types leads to near-certain receptor internalization. As the unbinding rate *k*_u_ increases, the fidelity also increases, as the receptor with lower unbinding rate *k*_u_ becomes internalized more frequently than the receptor with higher unbinding rate 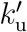. The fidelity in Eq. 4 saturates at *A*^2^ = 10^2^ as the unbinding rate becomes high, corresponding to the internalization probability for both ligands described by the limiting power law in Fig. 2A. The discrimination between different ligands, as quantified by the fidelity, is bounded from above by 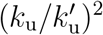.

With *k*_u,EGF_ = 0.05 s^−1^ and *k*_u,EREG_ = 0.5 s^−1^, the fidelity of EGF binding and internalization relative to epiregulin is approximately 20 (black circle in Fig. 2B), well below the saturated fidelity of 100. Receptors bound to EGF are approximately 20× more likely to be internalized than receptors bound to epiregulin. This fidelity range leaves ligand internalization sensitive to changes in EGFR dynamics, as illustrated in Fig. 2B. Adjusting the rate of clathrin recruitment *k*_c_ or internalization *k*_i_ will modestly change fidelity, with an order of magnitude decrease in either of these parameters nearly doubling the fidelity to approximately 35 and an order of magnitude increase in either of these parameters reducing the fidelity by almost two-thirds to approximately 8. The fidelity is less sensitive to the clathrin domain exit rate of receptors, *k*_-c_, with an order of magnitude increase in *k*_-c_ reducing the fidelity by approximately one-quarter, and an order of magnitude decrease in *k*_-c_ leaving the fidelity essentially unchanged.

Fidelity = *A*, the level at which fidelity matches the unbinding rate ratio, for 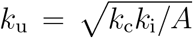. Thus the clathrin entry rate *k*_c_, internalization rate *k*_i_, and ratio of unbinding rates *A* control the unbinding rate at which the fidelity exceeds the unbinding rate ratio.

Figures 2C,D show the tradeoff between EGFR ligand fidelity and internalization probability, similar to previously noted tradeoffs [38, 39, 40, 41, 42]. The low fidelity limit for a given internalization probability occurs for a strongly two-stage and single-clathrin-entry process (*k*_i_, *k*_c_ ≫ *k*_-c_; *k*_c_ = *k*_i_),

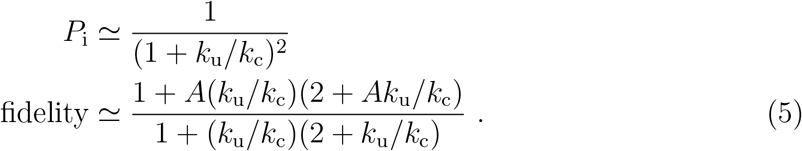

For the low fidelity limit, the internalization probability *P*_i_ and fidelity are controlled by a single parameter combination *k*_u_*/k*_c_ once the unbinding rate ratio *A* is selected. The high fidelity limit for a given internalization probability occurs for multiple cycles into and out of clathrin and high internalization (*k*_c_, *k*_i_ ≫ *k*_u_; *k*_-c_ ≫ *k*_c_, *k*_i_),

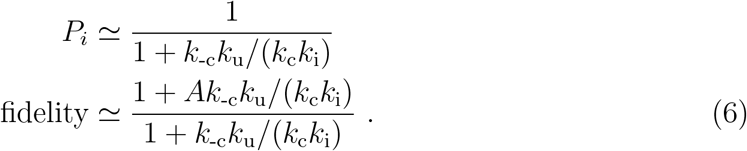

For the high fidelity limit, the internalization probability *P*_i_ and fidelity are controlled by a single parameter combination *k*_-c_*k*_u_*/*(*k*_c_*k*_i_) once the unbinding rate ratio *A* is selected. The high fidelity limit indicates that for fidelity to increase, the receptor internalization probability must decrease, as some EGFR must fail to internalize for stimulation by different ligands to become substantially distinct. A high fidelity for a given internalization probability may be desirable, as it provides a large distinction between the signaling activity due to the two ligands for a given number of receptors internalized. Figure 2C shows that the estimated EGFR parameters for EGF vs EREG binding are near but meaningfully below the high fidelity limit, and that reducing the clathrin exit rate *k*_-c_ appears to push the fidelity quite close to the high fidelity limit.

### 2.3. Clathrin domain entry

The internalization probability and the fidelity arise from considering receptor visits to clathrin domains. The probability of zero clathrin visits is *P*_v_(0) = *k*_u_*/*(*k*_u_ + *k*_c_) while the probability of *n* ≥ 1 clathrin visits is

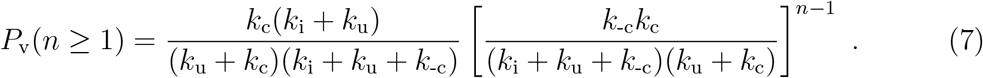

EGF-bound receptors average 0.73 visits to a clathrin domain and epiregulin-bound receptors average 0.16 visits. While the number of visits to clathrin domains before ligand unbinding or internalization are substantially affected by variation in EGFR dynamics for low unbinding rate *k*_u_ (Fig. 3A), there is more limited parameter sensitivity in the neighborhood of the EGF unbinding rate *k*_u,EGF_ = 0.05 s^−1^ (black circle in Fig. 3A) and the epiregulin unbinding rate *k*_u,EGF_ = 0.5 s^−1^ (orange circle in Fig. 3A). In this neighborhood, the clathrin recruitment rate *k*_c_ meaningfully impacts the number of visits to a clathrin domain, with an order of magnitude increase in *k*_c_ increasing the number of clathrin domain visits by 1.85× (EGF) and 4.4× (EREG) and an order of magnitude decrease in *k*_c_ decreasing the number of clathrin domain visits by 5.9× (EGF) and 8.7× (EREG).

**Figure 3.**
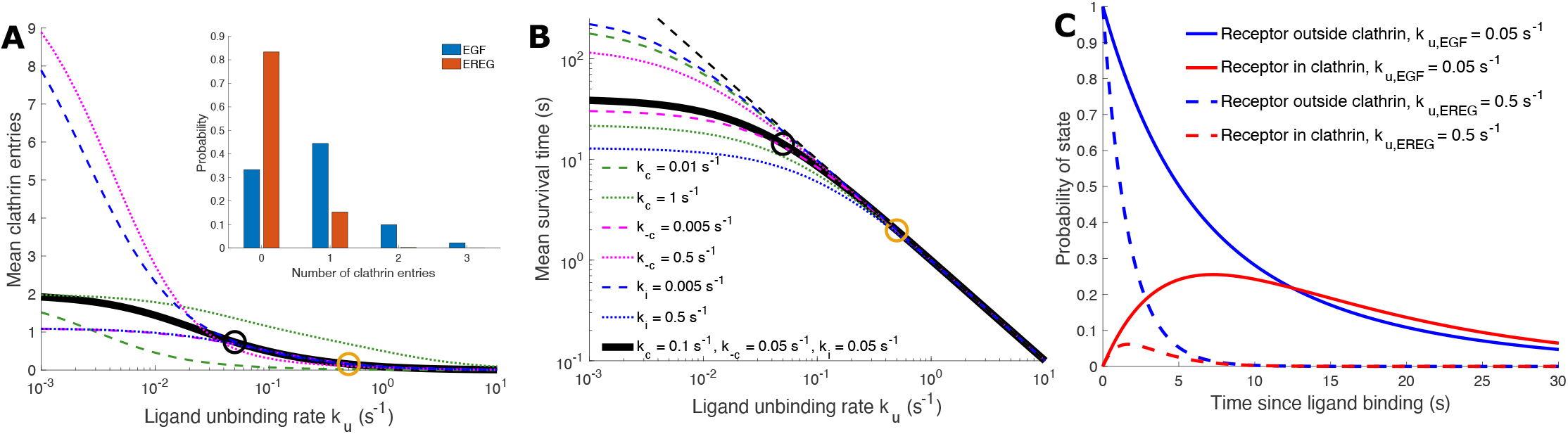
EGFR dynamics with kinetic model. (A) Mean number of entries into a clathrin domain as ligand unbinding rate *k*_u_ is varied. Curves as indicated by legend in (B); thick solid black curve for estimated EGFR parameters (see text), and other curves with one parameter value changed compared to thick black curve, as indicated. Black circle shows mean clathrin entries for *k*_u,EGF_ = 0.05 s^−1^ and orange circle for *k*_u,EREG_ = 0.5 s^−1^. Inset of (A) shows the distribution of the number of clathrin entries for the thick black curve parameters for *k*_u,EGF_ and *k*_u,EREG_. (B) Mean survival time (i.e. time from ligand binding until either internalization or ligand unbinding) as the unbinding rate *k*_u_ is varied. Black circle for *k*_u,EGF_ = 0.05 s^−1^ and orange circle for *k*_u,EREG_ = 0.5 s^−1^. Black dashed line is high *k*_u_ limiting behavior of mean survival time ≃ 1*/k*_u_. (C) Probability of receptor being within or outside a clathrin domain as time since ligand binding increases, for *k*_u,EGF_ and *k*_u,EREG_.

The inset of Fig. 3A shows the fraction of receptors undergoing zero, one, two, and three clathrin domain visits, for EGF and epiregulin unbinding rates. For both EGF and epiregulin, it is rare for a receptor to undergo more than three distinct visits to a clathrin domain before receptor internalization or ligand unbinding. The inset of Fig. 3A also shows that EGF-bound receptors mostly visit clathrin domains zero times, once, or twice, with almost half visiting once; while epiregulin-bound receptors mostly visit clathrin domains zero times or once, with more than 80% not visiting clathrin domains.

Varying unbinding rate also impacts the time before either receptor internalization or ligand unbinding, shown as the mean survival time in Fig. 3B. Similar to the internalization probability (Fig. 2A), the mean survival time is flatter for low unbinding rate *k*_u_ and then decreases before reaching a limiting power law for large *k*_u_. EGF-bound receptors have a mean survival time of approximately 14 s (Fig. 3B, black circle) and epiregulin-bound receptors have a mean survival time of approximately 2 s (orange circle). While at high *k*_u_ the internalization probability (Fig. 2A) depends on the parameters describing EGFR dynamics and at low *k*_u_ the internalization probability converges independent of parameters (converging because *P*_i_ cannot exceed one), the mean survival time (Fig. 3B) depends on parameters at low *k*_u_ and converges independent of parameters for high *k*_u_. This convergence at high *k*_u_ occurs because unbinding becomes dominant over internalization, and the mean survival time becomes approximately 1*/k*_u_.

Figure 3C shows the time evolution following ligand binding of the occupation probability of ligand-bound (*R*_L_) and clathrin-localized (*R*_C_) receptor states (see Appendix for equations), for both EGF and epiregulin binding. Consistent with the faster unbinding and lower survival time for epiregulin-bound receptors compared to EGF-bound receptors, the probability of occupying either state decreases rapidly for epiregulin-bound receptors. The peak in clathrin-localized epiregulin-bound receptors reaches approximately 6% of receptors at 1 – 2 s, and decays to below 1% after 8 s. In contrast, many more EGF-bound receptors become and remain localized to clathrin domains, with a peak of approximately 26% of receptors occurring at 7 – 8 s, and decaying to below 1% after more than 50 s.

### 2.4. Spatial model

In addition to behavior of the kinetic model of EGFR clathrin association, internalization, and ligand unbinding described above, we explore a two-dimensional spatial model (Fig. 1B). Receptors begin with a ligand bound, diffusing with *D* = 0.2 *μ*m^2^*/*s to encounter randomly distributed (but not overlapping) circular clathrin domains. Energy barriers limit entry into clathrin domains, from a receptor attempting to diffuse onto a clathrin domain site from outside; and exit from clathrin domains, from a receptor attempting to diffuse onto a site outside a clathrin domain from a site inside. Receptors diffuse and enter and exit clathrin domains until the receptor is internalized within a clathrin domain or the ligand unbinds.

Experiments suggest that EGFR can become localized to a clathrin domain either through induction or stabilization of a nascent clathrin domain, or by recruitment to an existing clathrin domain [14, 15, 16, 17]. We treat both of these cases as a diffusive encounter with a circular clathrin domain, as EGFR induction or stabilization of a nascent clathrin domain will involve diffusive search by the receptor for clathrin domain components. For diffusive search by EGFR for an existing domain, which most closely matches this spatial model, the domain entry energy barrier is adjusted to match the clathrin entry rate estimated above (see next section). EGFR exit from a clathrin domain is often due to clathrin domain disassembly [48, 57, 58]. Accordingly, when an EGFR exits a clathrin domain in the model (crossing an energy barrier with height adjusted to match the clathrin exit rate estimated above; see next section) the clathrin domain is moved to another location. See Appendix for further simulation details.

#### 2.4.1. Spatial model parameters

While the internalization rate *k*_i_ and ligand unbinding rate *k*_u_ in the spatial model and simulation directly correspond to these same parameters in the kinetic model, the clathrin domain entry rate *k*_c_ and exit rate *k*_-c_ in the kinetic model do not have direct correspondence in the spatial model. With the spatial model, *k*_c_ is controlled by the clathrin domain entry energy barrier Δ*E*_enter_, clathrin domain radius *r*_c_, clathrin domain concentration *c*_c_, and receptor diffusivity *D. k*_-c_ is controlled by the clathrin domain exit energy barrier Δ*E*_exit_, clathrin domain radius *r*_c_, and receptor diffusivity *D*.

The effective entry rate into the clathrin domain *k*_c_ from the kinetic model is represented in the spatial model by diffusive search for the clathrin domains followed by crossing the entry energy barrier Δ*E*_enter_. Diffusive search is determined by the clathrin domain radius *r*_c_, clathrin domain concentration *c*_c_, and receptor diffusivity *D*. By assuming that the time to enter the clathrin domain is the sum of an equilibration time period for even distribution on the plasma membrane and a time period for equilibrated receptors that are adjacent to the clathrin domain to overcome the entry energy barrier, we estimate (see Appendix for derivation details) the mean time to enter as

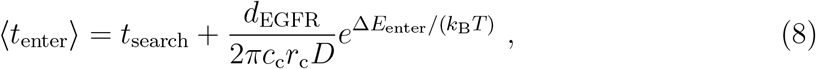

with *d*_EGFR_ = 10 nm the EGFR diameter. *k*_B_ is Boltzmann’s constant and *T* is absolute temperature, together representing a thermal energy scale.

In the simulation, changing Δ*E*_enter_ in turn changes the mean time to enter a clathrin domain, shown in Fig. 4A. While not an exact match, the ⟨*t*_enter_⟩ estimate (Eq. 8) has a similar form and magnitude to the simulation domain entry times. Δ*E*_enter_ = 0 corresponds to diffusion-limited entry of receptors into clathrin domains, where any receptor that reaches a clathrin domain will enter. A mean clathrin domain entry time of 1.47 s (first entry of each trajectory) and 1.71 s (all entries of each trajectory) occurs for Δ*E*_enter_ = 0, the diffusion-limited rate. This is larger than the diffusion-limited search time estimated above for the kinetic model of ≈ 1 s, showing the random distribution of clathrin domains in the spatial model leads to somewhat larger diffusion-limited search times. For low Δ*E*_enter_ ≲ 3 *k*_B_*T* the time to enter a clathrin domain is largely the diffusive encounter time (first term in Eq. 8) as there is a high chance of a receptor entering an encountered clathrin domain. For high Δ*E*_enter_ ≳ 4 *k*_B_*T*, the time to enter a clathrin domain is largely controlled by the time spent near clathrin domains over time and the difficulty of entering a clathrin domain — for Fig. 4A, this is indicated by the simulated mean time to enter a clathrin domain 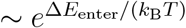 (second term in Eq. 8).

**Figure 4.**
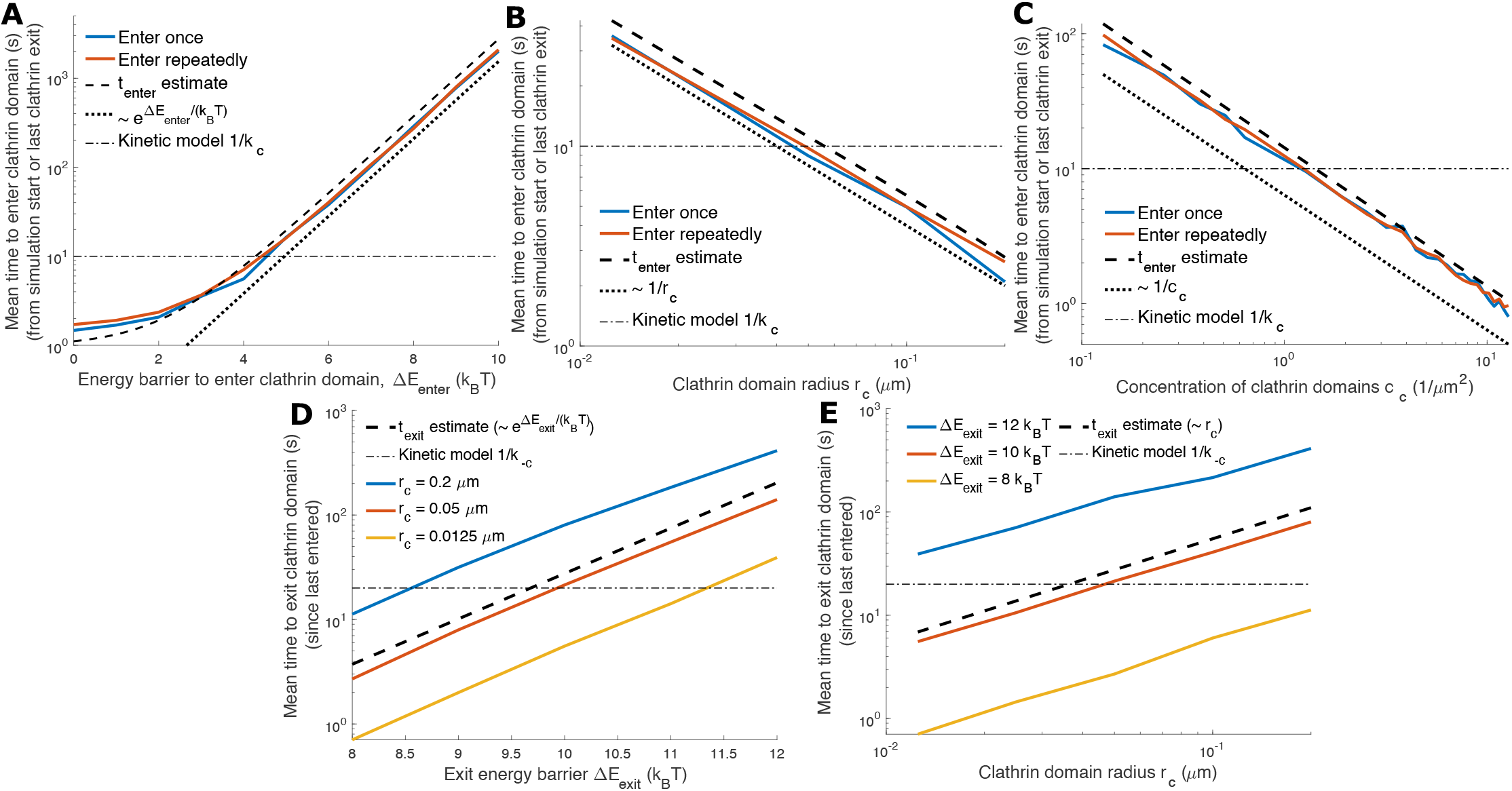
Spatial model parameter variation. (A,B,C) the mean time for a receptor to enter a clathrin domain as (A) the energy barrier to enter a clathrin domain Δ*E*_enter_, (B) the clathrin domain radius *r*_c_, and (C) the concentration of clathrin domains *c*_c_ are varied. Blue solid curves are the mean time for receptors initially in a random position outside a clathrin domain to first enter a clathrin domain. Red solid curves are the mean time for multiple clathrin entries including the initial entry, with subsequent entries contributing to the mean as the time interval from a clathrin domain exit to the next entry. Black dashed curves are the clathrin domain entry time estimate from Eq. 8. The dotted black curve is proportional to (A) the exponential 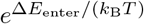, (B) the inverse of clathrin domain radius *r*_c_, and (C) the inverse of clathrin domain concentration *c*_c_. The dashed-dotted black curve is the mean time estimated from experimental data of 1*/k*_c_ = 10 s. *k*_u_ = 10^−5^ s^−1^ to allow time for multiple entries and Δ*E*_exit_ = 4.5 *k*_B_*T* to allow more efficient simulations. Means averaged over 100 runs. (D,E) Mean time for a receptor in a clathrin domain to exit the clathrin domain as the energy barrier to exit Δ*E*_exit_ and (E) the clathrin domain radius *r*_c_ are varied. Solid curves in (D) are different clathrin domain radii *r*_c_ and in (E) are different clathrin exit energy barriers Δ*E*_exit_, as indicated. Black dashed curves are the clathrin domain exit time estimate from Eq. 9 which is proportional to 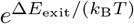 and *r*_c_. The dashed-dotted black curves are the mean time estimated from experimental data of 1*/k*_-c_ = 20 s. *k*_u_ = 10^−5^ s^−1^ to allow more clathrin domain exits and Δ*E*_enter_ = 0 for more efficient simulations. Means averaged over 100 runs.

In Fig. 4A, Δ*E*_enter_ ≃ 4.4 *k*_B_*T* leads to a clathrin domain enter time of 10 s (corresponding to *k*_c_ = 0.1 s^−1^ from the kinetic model). We will use Δ*E*_enter_ = 4.4 *k*_B_*T* hereafter, unless otherwise stated.

Figure 4B shows that the simulated mean time to enter a clathrin domain is inversely proportional to the domain radius, ∼ 1*/r*_c_, as expected from Eq. 8 for larger Δ*E*_enter_. Figure 4C shows the simulated mean time to enter a clathrin domain is inversely proportional to the clathrin domain concentration, ∼ 1*/c*_c_, as expected from Eq. 8 for larger Δ*E*_enter_. In Fig. 4B,C the Eq. 8 prediction for the time to enter a clathrin domain has the same functional dependence on *r*_c_ and *c*_c_ as the simulation results, but exceeds the simulation results by up to 20 – 30%, largely describing clathrin domain entry with no free parameters. While we do not use the data shown in Fig. 4B,C to determine *r*_c_ and *c*_c_ values, the figures indicate how the clathrin domain entry time depends on these parameters.

The effective exit rate from clathrin domains, *k*_-c_ from the kinetic model, is represented in the spatial model by diffusive search for the clathrin domain boundary from within followed by crossing the exit energy barrier Δ*E*_exit_. Diffusive search is determined by the clathrin domain radius *r*_c_ and receptor diffusivity *D*. The energy barrier Δ*E*_exit_, together with the diffusive search parameters, determine *k*_-c_ for the spatial model.

The kinetic model, with parameters estimated from experimental measurements, has a rate of exiting clathrin domains *k*_-c_ = 0.05 s^−1^. For free diffusion in two dimensions, ⟨*r*^2^⟩ = 4*Dt*, so for a clathrin domain radius *r*_c_ = 0.05 *μ*m and EGF receptor diffusivity *D* = 0.2 *μ*m^2^*/*s, the typical time to reach the clathrin boundary from within is ≲ 0.01 s, which is much shorter than the typical 20 s to exit a clathrin domain. By assuming that the receptor location has equilibrated within the domain and the exit time is for the receptors adjacent to the clathrin domain boundary to overcome the exit energy barrier, we estimate (see Appendix for details) the mean time to exit as

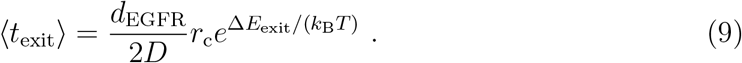

In the simulation of the spatial model, increasing Δ*E*_exit_ (Fig. 4D) and increasing the clathrin domain radius *r*_c_ (Fig. 4E) each increase the mean time to exit a clathrin domain. While the clathrin domain exit time estimate (Eq. 9) does not exactly match the spatial model simulation (exceeding the simulation by up to 45%), the estimate is of similar order of magnitude and matches the simulation data trend of mean clathrin domain exit time proportionality to exit barrier Boltzmann factor 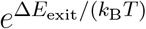 and clathrin domain radius *r*_c_.

Figure 4D shows that for a clathrin domain radius *r*_c_ = 0.05 *μ*m, as estimated above, the kinetic model clathrin domain exit rate *k*_-c_ = 0.05 s^−1^ is achieved for an exit barrier Δ*E* ≃ 10 *k*_B_*T* . We will use Δ*E*_exit_ = 10 *k*_B_*T* hereafter unless otherwise stated.

#### 2.4.2. Clathrin domain entry

We now examine in more detail the entry of receptors into clathrin domains. Figure 5A shows the probability density of times *t* to enter a clathrin domain, for receptors with ligands that do not unbind (*k*_u_ → 0). The time distributions appear approximately exponential at most times, ∼ exp(−*t/τ*), consistent with previous diffusive search and reaction time densities [59]. Adding the clathrin domain entry energy barrier of 4.4 *k*_B_*T* increases the time constant *τ* by approximately a factor of five.

**Figure 5.**
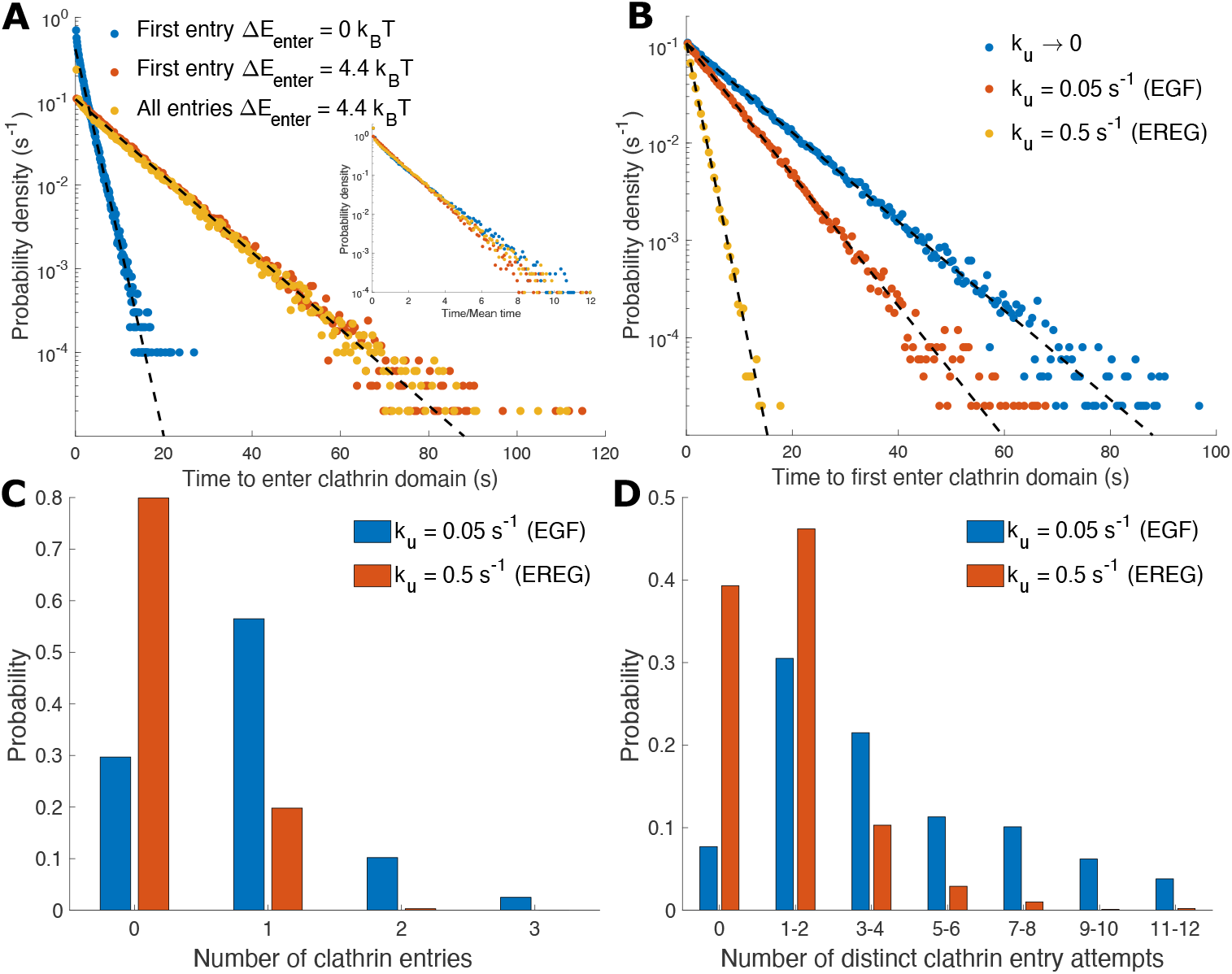
EGFR entry into clathrin domains for spatial model. (A) Probability density of times for receptor to enter a clathrin domain. First entry is for the initial receptor entry without subsequent entries. All entries is for multiple clathrin domain entries including the initial entry. *k*_u_ → 0 such that receptors enter and exit clathrin domains until internalized. Entry type and clathrin domain entry energy barrier Δ*E*_enter_ as indicated for data points. Black dashed lines are ∼ *e*^*−t/τ*^ with *τ* = 1.88 s (through blue) and *τ* = 9.5 s (through red/yellow). Mean time to enter is 1.56 s (blue), 9.5 s (red), and 8.76 s (yellow). Inset of (A) is the data of the main plot with times scaled by the mean time for each entry type and clathrin entry energy barrier. 10^5^ clathrin domain entries for each distribution. (B) Probability density of times for receptor to first enter a clathrin domain for different ligand unbinding rates *k*_u_. Mean time to first enter is 9.5 s (blue), 6.4 s (red), and 1.6 s (yellow). 32% (*k*_u,EGF_ = 0.05 s^−1^), and 82% (*k*_u,EREG_ = 0.5 s^−1^) of trajectories are not included in the distribution as they do not enter a clathrin domain prior to ligand unbinding, from 10^5^ trajectories for each of the three *k*_u_. Black dashed lines are 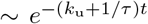 with *τ* = 9.5 s. Δ*E*_enter_ = 4.4 *k*_B_*T* . (C) Distribution of the number of clathrin entries and (D) distribution of the number of distinct clathrin entry attempts for *k*_u,EGF_ and *k*_u,EREG_. An entry attempt is distinct if the attempted entry is for a different clathrin domain than the previous attempt or if the receptor entered a clathrin domain between attempts. The mean number of clathrin entries is 0.9 (*k*_u,EGF_ = 0.05 s^−1^) and 0.2 (*k*_u,EREG_ = 0.5 s^−1^) . The mean number of distinct clathrin entry attempts is 5.1 (*k*_u,EGF_ = 0.05 s^−1^) and 1.2 (*k*_u,EREG_ = 0.5 s^−1^). (C) and (D) are from 1000 runs.

The time distribution to enter a clathrin domain on the first instance (after starting at a random position outside of clathrin domains) is very similar to the distribution to enter after exiting another clathrin domain. With clathrin domains moving after a receptor exits (see spatial model description in Appendix), the random initial position and clathrin domain exit position appear to have sufficiently similar distributions to lead to the similar clathrin entry time distributions. As all three distributions are exponential, scaling the first entry distributions with and without an energy barrier, and the repeated entry distribution, by their mean times leads to collapse of the clathrin entry time distributions (Fig. 5A inset).

Figure 5B examines the effect of nonzero unbinding rate *k*_u_ on the time distribution to first enter a clathrin domain with Δ*E*_enter_ = 4.4 *k*_B_*T* (expected to be similar to the distribution for repeated entries, as for Fig. 5A), with the probability density normalized to the total number of trajectories, including the receptors with trajectories that end with ligands that unbind prior to clathrin domain entry. These clathrin entry distributions are exponential in time, ∼ exp[−(*k*_u_ + 1*/τ*)*t*], with *τ* from Fig. 5A.

For unbinding rates corresponding to EGF (*k*_u,EGF_ = 0.05 s^−1^) and epiregulin (*k*_u,EREG_ = 0.5 s^−1^), Fig. 5C shows the number of clathrin domains that a receptor enters before ligand unbinding or internalization. The number of clathrin entries is very similar to those in the inset of Fig. 3A for the kinetic model, with EGF-bound receptors mostly visiting a clathrin domain zero times, once, or twice; and epiregulin-bound receptors mostly visiting clathrin domains zero times or once.

We also examined the number of distinct clathrin domains that a receptor attempts to enter, as a representation of how many clathrin domains a receptor will encounter, which will be related to the length of the diffusive search before a receptor successfully enters a clathrin domain. An entry attempt is considered distinct if for a different clathrin domain than the previous attempt, or if the receptor successfully entered a clathrin domain between attempts. EGF-bound receptors undergo a mean of approximately 5.1 clathrin entry attempts before ligand unbinding or internalization, while epiregulin-bound receptors undergo approximately 1.2 clathrin entry attempts. Figure 5D shows the distribution of the number of attempts. EGF-bound receptors rarely (*<* 10%) have EGF unbind prior to attempting clathrin domain entry, with approximately 30% of EGF-bound receptors attempting entry to 1 – 2 clathrin domains, a further 30% attempting entry to 3 – 6 clathrin domains, and nearly 30% attempting entry to ≥ 7 clathrin domains. In contrast, approximately 40% of epiregulin-bound receptors have epiregulin unbind prior to attempting to enter a clathrin domain, with nearly half of epiregulin-bound receptors attempting entry to 1 – 2 clathrin domains, approximately 10% attempting entry to 3 – 4 clathrin domains, and *<* 5% attempting entry to ≥ 5 clathrin domains.

#### 2.4.3. Proofreading

We now explore proofreading with the spatial model. Specifically, we compare the internalization probability *P*_i_ vs ligand unbinding rate *k*_u_ relationship (as in Fig. 2A) for the spatial model to the kinetic model with the same parameter values. If this relationship is similar for the spatial and kinetic models, then the fidelity-unbinding rate relationship will be similar and there will be similar proofreading behavior. Overall, we find that the internalization probabilities for the kinetic and spatial models are nearly identical (Fig. 6), indicating that the kinetic proofreading behavior discussed above for the kinetic model is also reflected in spatial model behavior.

**Figure 6.**
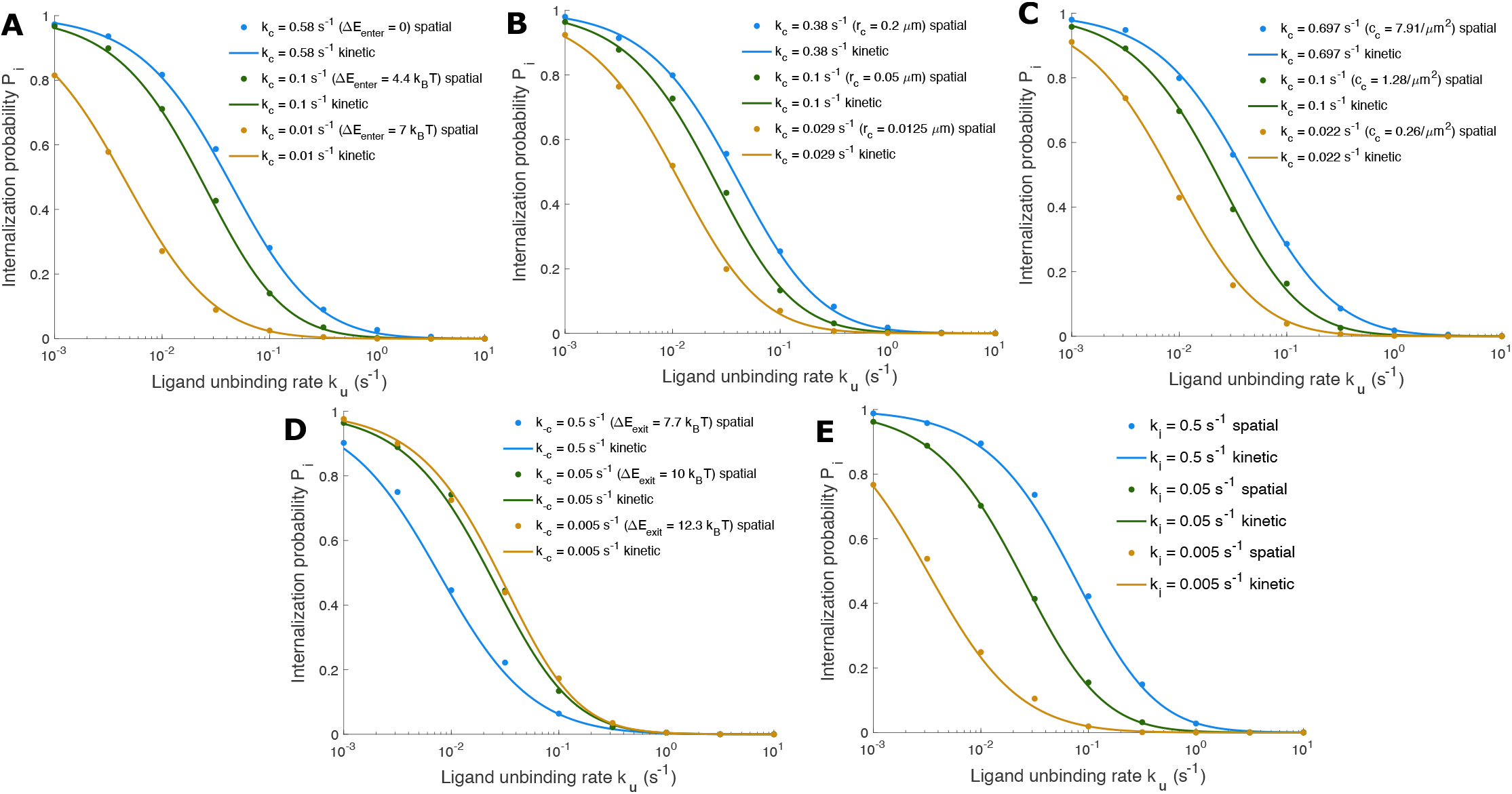
Internalization probabilities for the spatial model. (A – E) Internalization probability as ligand unbinding rate *k*_u_ is varied. Closed circles are from spatial model and solid curves are from kinetic model (Eq. 1) with equivalent parameter values. (A) Clathrin entry energy barrier Δ*E*_enter_ is varied to change *k*_c_ according to Fig. 4A, with *k*_c_ = 0.58 s^−1^ using Δ*E*_enter_ = 0, *k*_c_ = 0.1 s^−1^ using Δ*E*_enter_ = 4.4 *k*_B_*T*, and *k*_c_ = 0.01 s^−1^ using Δ*E*_enter_ = 7 *k*_B_*T* . (B) Clathrin domain radius *r*_c_ is varied to change *k*_c_ according to Fig. 4B, with *k*_c_ = 0.38 s^−1^ using *r*_c_ = 0.2 *μ*m, *k*_c_ = 0.1 s^−1^ using *r*_c_ = 0.05 *μ*m, and *k*_c_ = 0.029 s^−1^ using *r*_c_ = 0.0125 *μ*m. (C) Clathrin domain concentration *c*_c_ is varied to change *k*_c_ according to Fig. 4C, with *k*_c_ = 0.697 s^−1^ using *c*_c_ = 7.91*/μ*m^2^, *k*_c_ = 0.1 s^−1^ using *c*_c_ = 1.28*/μ*m^2^, and *k*_c_ = 0.022 s^−1^ using *c*_c_ = 0.26*/μ*m^2^. (D) Clathrin exit energy barrier Δ*E*_exit_ is varied to change *k*_-c_ according to Fig. 4D, with *k*_-c_ = 0.5 s^−1^ using Δ*E*_exit_ = 7.7 *k*_B_*T, k*_-c_ = 0.05 s^−1^ using Δ*E*_exit_ = 10 *k*_B_*T*, and *k*_-c_ = 0.005 s^−1^ using Δ*E*_exit_ = 12.3 *k*_B_*T* . (E) Internalization rate *k*_i_ is directly varied. Unless stated otherwise, parameters are Δ*E*_enter_ = 4.4 *k*_B_*T, r*_c_ = 0.05 *μ*m, and *c*_c_ = 1.28*/μ*m^2^, together corresponding to *k*_c_ = 0.1; Δ*E*_exit_ = 10 *k*_B_*T* corresponding to *k*_-c_ = 0.05 s^−1^; and *k*_i_ = 0.05 s^−1^. Spatial model results for all panels averaged over 1000 runs.

In Fig. 6A the clathrin entry energy barrier Δ*E*_enter_ is adjusted to vary the clathrin domain entry rate *k*_c_. For the spatial model, *k*_c_ cannot exceed approximately 0.58 s^−1^, because this is the diffusion-limited limit for clathrin domain encounter, as the shortest mean time to enter a clathrin domain for Δ*E*_enter_ = 0 is approximately 1*/*(0.58 s^−1^) ≃ 1.7 s. In Fig. 6B, *k*_c_ is varied by adjusting the clathrin domain radius *r*_c_; and in Fig. 6C, *k*_c_ is varied by adjusting the clathrin domain concentration *c*_c_. The adjustment of *k*_c_ via Δ*E*_enter_, *r*_c_, and *c*_c_ applies calibration from Figs. 4A–C.

In Fig. 6D the clathrin exit barrier Δ*E*_exit_ is adjusted to vary the clathrin domain exit rate *k*_-c_. The adjustment of *k*_-c_ via Δ*E*_exit_ applies calibration from Fig. 4D. In Fig. 6E receptor internalization rate in clathrin domains *k*_i_ is directly adjusted, without any calibration needed.

## 3. Discussion

We have investigated, using quantitative physical modeling, epidermal growth factor receptor (EGFR) internalization differences when stimulated by ligands with different receptor binding lifetimes. Discrimination of EGFR internalization modestly enhances underlying differences in ligand binding lifetimes. The results of the two models we develop, which we term kinetic and spatial models, are in agreement. Discrimination in EGFR internalization following ligand binding is a form of proofreading, combining the well-established kinetic proofreading [38, 39, 40, 41] with the more recently proposed spatial proofreading [42]. Different EGFR ligands lead to distinct cellular responses [28, 29, 30, 31] and EGFR signaling from endosomes following receptor internalization is distinct from EGFR signaling from the plasma membrane [18, 19, 20, 22, 23, 24]. This work shows that the internalization pathway of EGFR favors internalization and thus endosome-based EGFR signaling for ligands with longer binding lifetimes beyond what is expected from ligand unbinding rate ratios alone.

EGFR internalization following ligand binding combines kinetic proofreading and spatial proofreading. The first stage of EGFR internalization proofreading is spatial, as EGFR diffusively searches for a clathrin domain while ligand unbinding can occur. The second stage is kinetic, as the EGFR within a clathrin domain can be internalized while competing with ligand unbinding and clathrin domain exit. Although the first stage involves a diffusive search for a clathrin domain, the first stage is not diffusion limited — our spatial modeling suggests that EGFR typically encounters several distinct clathrin domains before successful recruitment into a clathrin domain, therefore often requiring a diffusive search comparable to or longer than the mean ligand binding lifetime.

The kinetic description of EGFR internalization proofreading (Fig. 1A) involves two stages, and is similar to the Hopfield description of two-stage kinetic proofreading between two substrates with an energy penalty for one substrate relative to the other [36, 40]. Initial substrate binding to form the first intermediate complex corresponds to ligand binding, the transition to the second intermediate complex from the first corresponds to recruitment to a clathrin domain, and the formation of the final product corresponds to internalization. Substrate unbinding from either intermediate state (generally assumed in the Hopfield description to have different rates from each complex) corresponds to ligand unbinding (which is described here with a single rate across states for EGFR), and returning to the first intermediate complex from the second corresponds to EGFR exiting a clathrin domain to diffuse on the plasma membrane. The Hopfield description energy penalty Δ*E* corresponds to the ratio of ligand unbinding rates 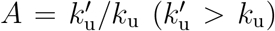, with *A* equivalent to the Boltzmann factor of the energy penalty 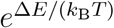. For Hopfield two-stage kinetic proofreading, the ratio of the final products between the two substrates cannot exceed 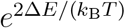 [40], and we equivalently find that EGFR internalization of long-relative to short-lifetime ligands cannot exceed *A*^2^.

There is a tradeoff between fidelity of internalization between two ligands and internalization probability, with a maximum fidelity for a given internalization probability, subject to the ratio of unbinding rates between the two ligands. This is similar to other performance tradeoffs in proofreading and other nonequilibrium processes, such as between speed and accuracy [60, 61]. We find that the fidelity of internalization between EGF and epiregulin binding approaches but is meaningfully below the maximum fidelity for a given internalization probability. While decreasing the rate at which EGFR exits clathrin domains can push the fidelity quite close to the upper fidelity limit, this may be a factor that is difficult for the cell to adjust, or constrained by how changes to clathrin domain dynamics may impact other processes.

Recent spatial proofreading work [42] considered a scenario where a complex with a bound substrate would undergo a diffusive search for a location facilitating catalysis, with an associated distribution of arrival times following substrate binding. Our spatial modeling suggests that EGFR typically undergoes multiple attempts to enter a clathrin domain before succeeding, instead following the exponential distribution of search times of a constant-rate or Poisson process.

EGFR internalization into endosomes facilitates important modes of signaling distinct from EGFR on the plasma membrane [18, 19, 20, 22, 23, 24], and ligand binding is important for this endosome-based EGFR signaling [25, 35]. For physiological EGFR parameters, and comparing EGF to epiregulin (EREG; approximately 10× faster unbinding than EGF) stimulation we find EGF binding is approximately 20× more likely to lead to EGFR internalization compared to EREG, modestly enhancing the unbinding rate difference.

However, the exact ratio between EGF- and EREG-stimulated internalization depends on the rates of recruitment to a clathrin domain, exiting a clathrin domain, and internalization. The receptor internalization fidelity is most sensitive to the clathrin recruitment and internalization rates, although a given fold-change in these rates is larger than the resulting internalization fidelity fold-change. Cells could adjust the factors behind these clathrin-domain rates to adjust discrimination in internalization of EGFR when stimulated by different ligands. Proteins involved in clathrin-mediated endocytosis can be perturbed or have their levels changed in cancer cells, altering clathrin-mediated endocytosis [62, 63, 64], suggesting that these changes in cancer cells may adjust the internalization and downstream signaling differences between EGFR ligands.

While we have emphasized signaling distinctions that may arise from differential internalization and endosomal localization of EGFR following binding of different ligand types, these internalization distinctions between ligand types also lead to differential depletion of EGFR from the plasma membrane, which is expected to have signaling consequences. With EGF binding inducing internalization with a substantially higher probability than EREG, EGF is expected to cause substantially more depletion of EGFR from the plasma membrane than EREG. Indeed, experiments show that EGF stimulation more rapidly depletes EGFR from the plasma membrane and depletes EGFR on the plasma membrane to a lower level, in comparison to EREG stimulation [25]. Although we focus on EGF and EREG stimulation, EGFR depletion from the plasma membrane following ligand stimulation varies across other ligands as well [25]. These depletion differences between ligand types, partially caused by different probabilities of internalization between ligand types, can then impact signaling persistence as EGFR must be located on the plasma membrane to respond to external ligand concentrations. Experiments show that EGF stimulation rapidly leads to EGFR signaling activity that decreases to less than half of the peak value after 20 minutes, while EREG and epigen (both lower affinity ligands) stimulation lead to longer-lived EGFR signaling activity persisting for an hour or longer [26]. The correspondence of stimulation by ligands with low receptor internalization probability to persistent signaling provides, in addition to endosomal localization, an additional mechanism to distinguish between EGFR stimulation by different ligands.

We examined two quantitative models of EGFR internalization, one entirely kinetic and another with spatial diffusion of EGFR to clathrin domains, and find that these two models are in close agreement. This agreement indicates that the time to enter a clathrin domain is distributed approximately exponentially, which is not a typical distribution for diffusion-controlled search times [65, 66]. However, our results suggest that EGFR recruitment to a clathrin domain is not a diffusion-limited process, and instead typically requires that EGFR visit several distinct clathrin domains. EGFR that are internalized after recruitment to a clathrin domain typically enter only one or two clathrin domains. This suggests that although EGFR can exit a clathrin domain (by exiting a persisting domain [67] or due to domain disassembly [48, 58]), EGFR does not typically exit clathrin domains repeatedly.

While recruitment of EGFR to a clathrin domain may involve receptor entry into an already-existing clathrin domain [16, 17], EGFR may also induce or stabilize the formation of clathrin domains [14, 15]. Our model describes recruitment to a clathrin domain either as a rate (kinetic model) or diffusive encounter with a clathrin domain followed by crossing an energy barrier (spatial model). Thus our spatial model does not explicitly account for EGFR recruitment to a clathrin domain via induction or stabilization of clathrin domain formation. However, as EGFR typically approaches several clathrin domains in the spatial model before entering a clathrin domain, this entry is well-approximated by a constant rate process, which is a closer match to recruitment via domain induction compared to a diffusion-limited search for the clathrin domain. Our spatial model also included escape from a clathrin domain as a diffusive search for the domain boundary followed by crossing an energy barrier. EGFR diffusivity inside a clathrin domain is likely limited due to interaction with domain components [8], and the spatial model description is aimed at producing relevant rates of clathrin escape. Through quantitative modeling, we have shown that the internalization of EGFR enhances unbinding rate differences between different EGFR ligands. This adds to other factors that allow EGFR to distinguish between ligands, such as differential stabilization of EGF receptor dimers [26] and differences in internalized receptor degradation and recycling to the plasma membrane [29]. This work also proposes that EGFR internalization discrimination between different stimulating ligands is an example of combined kinetic and spatial proofreading.

## Acknowledgments

This work was supported by a Toronto Metropolitan University Undergraduate Research Opportunity Award (J.A.L.), Natural Sciences and Engineering Research Council of Canada (NSERC) Discovery Grant (A.I.B.), start-up funds provided by the Toronto Metropolitan University Faculty of Science (A.I.B.), and a Canadian Institutes of Health Research (CIHR) Project Grant (C.N.A.).

## 4. Appendix

### 4.1. Kinetic model

#### 4.1.1. Calculating internalization probability

The probability that a receptor in state *R*_L_ enters a clathrin domain before ligand unbinding is *k*_c_*/*(*k*_c_ + *k*_u_). The probability that a receptor in state *R*_C_ is internalized before ligand unbinding is *k*_i_*/*(*k*_i_ +*k*_-c_ +*k*_u_) and the probability that a receptor in a clathrin domain returns to state *R*_L_ without ligand unbinding is *k*_-c_*/*(*k*_i_ + *k*_-c_ + *k*_u_). The receptor can transition back and forth between *R*_L_ and *R*_C_ before ligand unbinding or internalization. The probability *P*_i_ of receptor internalization is

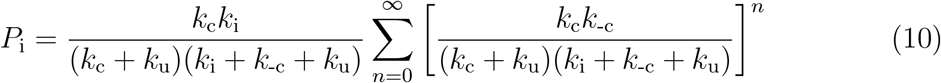

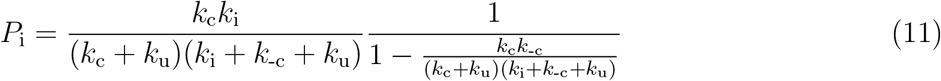

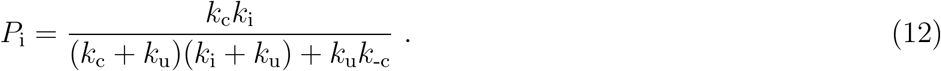

#### 4.1.2. Parameter estimation

We estimate the parameters *k*_c_, *k*_-c_, *k*_i_, and *k*_u_ for the kinetic model of EGFR dynamics. While the formation of clathrin domains can lead to the formation of clathrin-coated pits and internalization via endocytosis of any contained receptors, clathrin domains can also transiently form and disperse without an endocytosis event [48]. Successful internalization events of clathrin domains are estimated to occur over a 24 – 28 second time period [48]. We accordingly estimate the internalization rate as *k*_i_ ≈ 0.05 s^−1^, representing a mean period until internalization of 20 seconds following EGFR recruitment to a clathrin domain. We similarly estimate the transition of a receptor from inside a clathrin domain to outside a clathrin domain at a rate *k*_-c_ ≈ 0.05 s^−1^. This rate represents the release of EGFR upon clathrin domain disassembly, or the escape of EGFR from a retained clathrin domain. The upper limit of the clathrin domain recruitment rate *k*_c_ can be estimated as the diffusion-limited arrival rate of EGFR to clathrin domains from other regions of the cell membrane. The mean search time for a particle of diffusivity *D* to find a smaller absorbing cylinder of radius *r*_1_ when confined within a larger reflecting cylinder of radius *r*_2_ *> r*_1_ is [43]

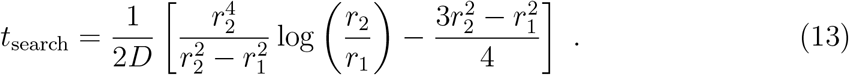

EGFR diffusivities have been experimentally measured as 0.08 - 0.20 *μ*m^2^*/*s [9], 0.2 *μ*m^2^*/*s [44, 45, 46], 0.25 *μ*m^2^*/*s [47], and 0.1 – 0.5 *μ*m^2^*/*s [8]. We use an intermediate value from these measurements, *D*_EGFR_ = 0.2 *μ*m^2^*/*s. Clathrin domains are approximately 100 nm in diameter [48], so the radius of the smaller absorbing circle is *r*_1_ ≈ 50 nm. Clathrin domains are approximately 1% of the membrane surface [48] (see Spatial model below), so with the rest of the circle representing the region nearest to a particular clathrin domain (if the receptor leaves the region nearest one clathrin domain, the receptor enters the region nearest to another clathrin domain), then *r*_2_ ≈ 500 nm. For *D*_EGFR_ ≈ 0.2 *μ*m^2^*/*s, *r*_2_ ≈ 500 nm, and *r*_1_ ≈ 50 nm, *t*_search_ ≈ 1 s. This sets a diffusion-limited upper bound on the rate clathrin domain entry of *k*_c,DL_ = 1*/t*_search_ ≈ 1 s^−1^. Recent experimental work [8] finds that approximately 50% of ligand-bound EGFR are confined to clathrin domains, or *R*_C_ ≈ *R*_L_. A steady-state concentration of clathrin-localized ligand-bound EGF receptors *R*_C,ss_ requires *k*_c_*R*_L,ss_ = (*k*_i_ + *k*_-c_)*R*_C,ss_. With *R*_C,ss_ = *R*_L,ss_, then *k*_c_ = *k*_i_ + *k*_-c_ ≈ 0.1 s^−1^. This is well under the diffusion-limited rate *k*_c,DL_ ≈ 1 s^−1^ and similar to the rate corresponding to a time period until the onset of slow diffusion following EGF stimulation of approximately 4 – 22 s [9].

#### 4.1.3. Time-dependence of R_L_ and R_C_ occupation

The time evolution of the probabilities *P*_L_ and *P*_C_, of respectively occupying the states *R*_L_ and *R*_C_, in the kinetic model shown in Fig. 1A is

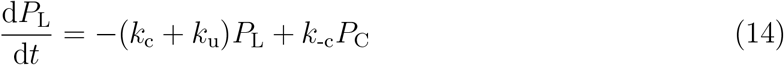

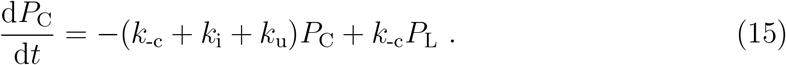

We write 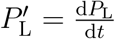 and 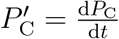. Taking the derivative of Eq. 14 with respect to time provides 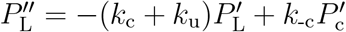 and then substituting Eq. 15 gives

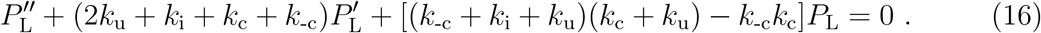

Equation 16 indicates a *P*_L_(*t*) of the form

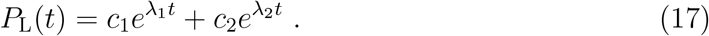

Equation 17 is substituted into Eq. 14 to find

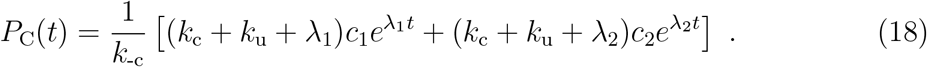

Following ligand binding EGFR is in state *R*_L_, so *P*_L_(*t* = 0) = 1 and *P*_C_(*t* = 0) = 0, leading from Eq. 18 to *c*_2_ = −[(*k*_c_ + *k*_u_ + *λ*_1_)*/*(*k*_c_ + *k*_u_ + *λ*_2_)]*c*_1_. The condition that *P*_L_(*t* = 0) = 1 then leads from Eq. 17 to *c*_1_ = (*k*_c_ + *k*_u_ + *λ*_2_)*/*(*λ*_1_ − *λ*_2_) and *c*_2_ = −(*k*_c_ + *k*_u_ + *λ*_1_)*/*(*λ*_2_ − *λ*_1_). Substituting Eq. 17 into Eq. 16 gives 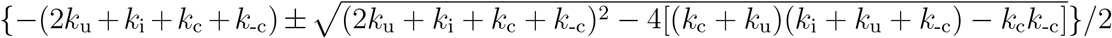.

### 4.2. Spatial model

#### 4.2.1. Simulation details

EGFR trajectories begin at a random position outside of clathrin domains with a ligand bound. Receptors diffuse on a two-dimensional square lattice with lattice site spacing Δ*x* = Δ*y* = 0.01 *μ*m. Experimental measurements suggest an individual EGFR diameter of approximately 7.2 nm [68], 10.6 nm [69], and 11.8 nm [70], and 11 nm between EGF receptor dimers [71], so we choose this lattice spacing equal to an estimated EGFR diameter *d*_EGFR_ = 10 nm. Receptors attempt a step to one of the four nearest-neighbor lattice sites every Δ*t* = 1.25 × 10^−4^ s, corresponding to a receptor diffusivity *D* = 0.2 *μ*m^2^*/*s. The simulated clathrin domains are circular with radius of *r*_c_ = 0.05 *μ*m (unless otherwise stated), corresponding to experimental measurements of clathrin domains approximately 100 nm in radius [48]. Ten clathrin domains (unless otherwise stated), are randomly placed in a square domain with periodic boundaries of side length 2.8 *μ*m without overlapping, corresponding to a concentration of 1.28*/μ*m^2^ and an area fraction of approximately 1%. This area fraction is calculated from measurements that the average lifetime of a clathrin domain is 46 s and the appearance per second of approximately three new clathrin clusters lasting at least 20 s per 10^8^ nm^2^ of membrane [48], which for 100 nm-diameter circular domains is (46 s)(2.4 × 10^4^ nm^2^*/*s)*/*10^8^ nm^2^ = 0.011 or approximately 1% of the membrane area occupied by a clathrin domain.

Attempted entries of receptors into a clathrin domain (i.e., attempted steps from a lattice site greater than distance *r*_c_ from the nearest clathrin domain center to a lattice site less than *r*_c_ from the domain center) are successful with probability 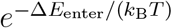, such that Δ*E*_enter_ represents an energy barrier [72] to entering clathrin domains. Attempted exit of receptors from a clathrin domain similarly encounters an energy barrier Δ*E*_exit_ while steps entirely outside or within a clathrin domain always succeed (no energy barrier). Exit of a receptor from a clathrin domain may correspond to disassembly of a clathrin domain [48, 58] or the exit from a persisting clathrin domain. We assume that both of these processes effectively represent a scenario where the clathrin domain has become unable to confine an EGFR, and thus move the clathrin domain to a new location upon receptor exit. The clathrin domain exit energy barrier Δ*E*_exit_ thus enforces a timescale before receptor exit from the domain.

Receptors inside a clathrin domain are internalized at a rate *k*_i_ (occurring with probability *k*_i_Δ*t* each timestep). Ligands unbind from any receptor with rate *k*_u_ (probability *k*_u_Δ*t*). Either receptor internalization or ligand unbinding ends the simulated trajectory of a receptor — if both internalization and unbinding are selected in the timestep, the process to occur is selected with probability proportional to the rate of each process.

#### 4.2.2. Estimating rate of entering clathrin domains

To estimate the mean time for a receptor to enter a clathrin domain, we sum the time period for a receptor to first diffusively encounter a clathrin domain and the time period to enter a clathrin domain after the first encounter, similar to splitting into diffusion and reaction timescales [59]. The time to first encounter a clathrin domain is estimated using *t*_search_ in Eq. 13. After this encounter it is assumed that the spatial distribution of the receptors relative to clathrin domains has equilibrated, and the fraction of time the receptor is adjacent to clathrin domains is equal to the area fraction of the membrane adjacent to clathrin domains, where the area within one EGFR diameter of a clathrin domain as a fraction of the total area is *f*_adjacent_ = *c*_c_(2*πr*_c_)*d*_EGFR_, with *c*_c_ the clathrin domain concentration. The rate of clathrin domain entry attempts from this adjacent region is *k*_attempt_ = 1*/*(4Δ*t*), where Δ*t* is the simulation timestep. Each attempt will be successful with a probablitiy 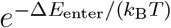. The estimated mean time to enter a clathrin domain is

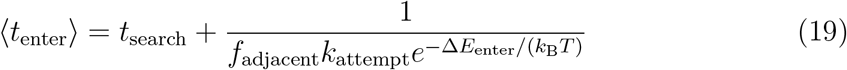

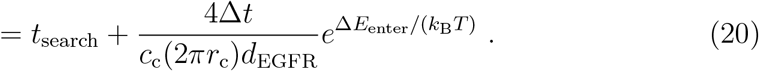

Diffusivity *D* = (Δ*x*)^2^*/*(4Δ*t*), with Δ*x* equal to the EGFR diameter *d*_EGFR_,

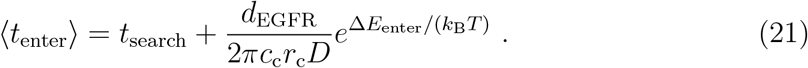

#### 4.2.3. Estimating rate of exiting clathrin domains

To estimate the mean time for a receptor to exit a clathrin domain, we assume that the receptor location has equilibrated within the domain and the exit time is for the receptors adjacent to the clathrin domain boundary to overcome the exit energy barrier, and the fraction of receptors near the domain boundary is equal to the fraction of the domain area adjacent to the domain boundary, or 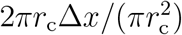. The rate of attempts to leave the domain when in this boundary-adjacent region is 1*/*(4Δ*t*), such that the average rate of attempting to leave the domain for clathrin-localized receptors is 2Δ*x/*(4Δ*tr*_c_). In two dimensions, diffusivity *D* = (Δ*x*)^2^*/*(4Δ*t*), so for Δ*x* = *d*_EGFR_, the attempt rate for leaving clathrin domains is 2*D/*(*r*_c_*d*_EGFR_). With clathrin domain exit energy barrier Δ*E*_exit_, the rate of leaving clathrin domains is 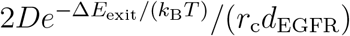 and the estimated mean time to exit a clathrin domain is

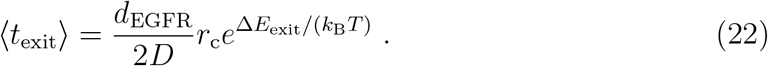

## Notes

### Competing Interest Statement

The authors have declared no competing interest.

## References

[1] Hernán E Grecco, Malte Schmick, and Philippe IH Bastiaens. Signaling from the living plasma membrane. Cell, 144(6):897–909, 2011.

[2] Yosef Yarden and Ben-Zion Shilo. Snapshot: EGFR signaling pathway. Cell, 131(5):1018–e1, 2007.

[3] Mark A Lemmon and Joseph Schlessinger. Cell signaling by receptor tyrosine kinases. Cell, 141(7):1117–1134, 2010.

[4] Ping Wee and Zhixiang Wang. Epidermal growth factor receptor cell proliferation signaling pathways. Cancers, 9(5):52, 2017.

[5] Sara Sigismund, Daniele Avanzato, and Letizia Lanzetti. Emerging functions of the EGFR in cancer. Molecular oncology, 12(1):3–20, 2018.

[6] Parthasarathy Seshacharyulu, Moorthy P Ponnusamy, Dhanya Haridas, Maneesh Jain, Apar K Ganti, and Surinder K Batra. Targeting the EGFR signaling pathway in cancer therapy. Expert opinion on therapeutic targets, 16(1):15–31, 2012.

[7] Curtis R Chong and Pasi A Jänne. The quest to overcome resistance to EGFR-targeted therapies in cancer. Nature medicine, 19(11):1389–1400, 2013.

[8] Michael G Sugiyama, Aidan I Brown, Jesus Vega-Lugo, Andrew M Scott, Khuloud Jaqaman, Gregory D Fairn, and Costin N Antonescu. Confinement of unliganded EGFR by tetraspanin nanodomains gates EGFR ligand binding and signaling. bioRxiv, pages 2022–03, 2022.

[9] Inhee Chung, Robert Akita, Richard Vandlen, Derek Toomre, Joseph Schlessinger, and Ira Mellman. Spatial control of EGF receptor activation by reversible dimerization on living cells. Nature, 464(7289):783–787, 2010.

[10] Sarah R Needham, Selene K Roberts, Anton Arkhipov, Venkatesh P Mysore, Christopher J Tynan, Laura C Zanetti-Domingues, Eric T Kim, Valeria Losasso, Dimitrios Korovesis, Michael Hirsch, et al. EGFR oligomerization organizes kinase-active dimers into competent signalling platforms. Nature communications, 7(1):13307, 2016.

[11] Lan K Nguyen, Walter Kolch, and Boris N Kholodenko. When ubiquitination meets phosphorylation: a systems biology perspective of EGFR/MAPK signalling. Cell Communication and Signaling, 11(1):1–15, 2013.

[12] Noga Kozer, Dipak Barua, Suzanne Orchard, Eduoard C Nice, Antony W Burgess, William S Hlavacek, and Andrew HA Clayton. Exploring higher-order EGFR oligomerisation and phosphorylation—a combined experimental and theoretical approach. Molecular BioSystems, 9(7):1849–1863, 2013.

[13] Lian Yi, Tujin Shi, Marina A Gritsenko, Chi-Yuet X’avia Chan, Thomas L Fillmore, Becky M Hess, Adam C Swensen, Tao Liu, Richard D Smith H Steven Wiley, et al. Targeted quantification of phosphorylation dynamics in the context of EGFR-MAPK pathway. Analytical chemistry, 90(8):5256–5263, 2018.

[14] Lene E Johannessen, Nina Marie Pedersen, Ketil Winther Pedersen, Inger Helene Madshus, and Espen Stang. Activation of the epidermal growth factor (EGF) receptor induces formation of EGF receptor-and Grb2-containing clathrin-coated pits. Molecular and cellular biology, 26(2):389–401, 2006.

[15] Yue Zhou and Hiroaki Sakurai. New trend in ligand-induced EGFR trafficking: A dual-mode clathrin-mediated endocytosis model. Journal of Proteomics, page 104503, 2022.

[16] Espen Stang, Frøydis D Blystad, Maja Kazazic, Vibeke Bertelsen, Tonje Brodahl, Camilla Raiborg, Harald Stenmark, and Inger Helene Madshus. Cbl-dependent ubiquitination is required for progression of EGF receptors into clathrin-coated pits. Molecular biology of the cell, 15(8):3591– 3604, 2004.

[17] Maja Kazazic, Vibeke Bertelsen, Ketil Winther Pedersen, Tram Thu Vuong, Michael Vibo Grandal, Marianne Skeie Rødland, Linton M Traub, Espen Stang, and Inger Helene Madshus. Epsin 1 is involved in recruitment of ubiquitinated EGF receptors into clathrin-coated pits. Traffic, 10(2):235–245, 2009.

[18] Sara Sigismund, Elisabetta Argenzio, Daniela Tosoni, Elena Cavallaro, Simona Polo, and Pier Paolo Di Fiore. Clathrin-mediated internalization is essential for sustained EGFR signaling but dispensable for degradation. Developmental cell, 15(2):209–219, 2008.

[19] Lasse Henriksen, Michael Vibo Grandal, Stine Louise Jeppe Knudsen, Bo van Deurs, and Lene Melsæther Grøvdal. Internalization mechanisms of the epidermal growth factor receptor after activation with different ligands. PloS one, 8(3):e58148, 2013.

[20] Mark von Zastrow and Alexander Sorkin. Signaling on the endocytic pathway. Current opinion in cell biology, 19(4):436–445, 2007.

[21] Alejandra Tomas, Clare E Futter, and Emily R Eden. EGF receptor trafficking: consequences for signaling and cancer. Trends in cell biology, 24(1):26–34, 2014.

[22] Amandio V Vieira, Christophe Lamaze, and Sandra L Schmid. Control of EGF receptor signaling by clathrin-mediated endocytosis. Science, 274(5295):2086–2089, 1996.

[23] Patrick Burke, Kevin Schooler, and H Steven Wiley. Regulation of epidermal growth factor receptor signaling by endocytosis and intracellular trafficking. Molecular biology of the cell, 12(6):1897–1910, 2001.

[24] Lukasz Sadowski, Iwona Pilecka, and Marta Miaczynska. Signaling from endosomes: location makes a difference. Experimental cell research, 315(9):1601–1609, 2009.

[25] Kirstine Roepstorff, Michael Vibo Grandal, Lasse Henriksen, Stine Louise Jeppe Knudsen, Mads Lerdrup, Lene Grøvdal, Berthe Marie Willumsen, and Bo Van Deurs. Differential effects of EGFR ligands on endocytic sorting of the receptor. Traffic, 10(8):1115–1127, 2009.

[26] Daniel M Freed, Nicholas J Bessman, Anatoly Kiyatkin, Emanuel Salazar-Cavazos, Patrick O Byrne, Jason O Moore, Christopher C Valley, Kathryn M Ferguson, Daniel J Leahy, Diane S Lidke, et al. EGFR ligands differentially stabilize receptor dimers to specify signaling kinetics. Cell, 171(3):683–695, 2017.

[27] Jennifer T Jones, Robert W Akita, and Mark X Sliwkowski. Binding specificities and affinities of egf domains for ErbB receptors. FEBS letters, 447(2-3):227–231, 1999.

[28] Kristy J Wilson, Jennifer L Gilmore, John Foley, Mark A Lemmon, and David J Riese II. Functional selectivity of EGF family peptide growth factors: implications for cancer. Pharmacology & therapeutics, 122(1):1–8, 2009.

[29] Stine Louise Jeppe Knudsen, Anni Sieu Wai Mac, Lasse Henriksen, Bo van Deurs, and Lene Melsæther Grøvdal. EGFR signaling patterns are regulated by its different ligands. Growth factors, 32(5):155–163, 2014.

[30] Carmen Berasain and Matías A Avila. Amphiregulin. In Seminars in cell & developmental biology, volume 28, pages 31–41. Elsevier, 2014.

[31] Tom Ronan, Jennifer L Macdonald-Obermann, Lorel Huelsmann, Nicholas J Bessman, Kristen M Naegle, and Linda J Pike. Different epidermal growth factor receptor (EGFR) agonists produce unique signatures for the recruitment of downstream signaling proteins. Journal of Biological Chemistry, 291(11):5528–5540, 2016.

[32] N Solic and DE Davies. Differential effects of EGF and amphiregulin on adhesion molecule expression and migration of colon carcinoma cells. Experimental cell research, 234(2):465–476, 1997.

[33] Eunkyung Chung, Ramona Graves-Deal, Jeffrey L Franklin, and Robert J Coffey. Differential effects of amphiregulin and TGF-α on the morphology of MDCK cells. Experimental cell research, 309(1):149–160, 2005.

[34] Jerusa AQA Faria, Carolina de Andrade, Alfredo M Goes, Michele A Rodrigues, and Dawidson A Gomes. Effects of different ligands on epidermal growth factor receptor (EGFR) nuclear translocation. Biochemical and biophysical research communications, 478(1):39–45, 2016.

[35] Anthony R French, Douglas K Tadaki, Salil K Niyogi, and Douglas A Lauffenburger. Intracellular trafficking of epidermal growth factor family ligands is directly influenced by the pH sensitivity of the receptor/ligand interaction. Journal of Biological Chemistry, 270(9):4334–4340, 1995.

[36] John J Hopfield. Kinetic proofreading: a new mechanism for reducing errors in biosynthetic processes requiring high specificity. Proceedings of the National Academy of Sciences, 71(10):4135–4139, 1974.

[37] Jacques Ninio. Kinetic amplification of enzyme discrimination. Biochimie, 57(5):587–595, 1975.

[38] Timothy W McKeithan. Kinetic proofreading in T-cell receptor signal transduction. Proceedings of the national academy of sciences, 92(11):5042–5046, 1995.

[39] Peter S Swain and Eric D Siggia. The role of proofreading in signal transduction specificity. Biophysical journal, 82(6):2928–2933, 2002.

[40] Arvind Murugan, David A Huse, and Stanislas Leibler. Speed, dissipation, and error in kinetic proofreading. Proceedings of the National Academy of Sciences, 109(30):12034–12039, 2012.

[41] Wenping Cui and Pankaj Mehta. Identifying feasible operating regimes for early T-cell recognition: The speed, energy, accuracy trade-off in kinetic proofreading and adaptive sorting. PloS one, 13(8):e0202331, 2018.

[42] Vahe Galstyan, Kabir Husain, Fangzhou Xiao, Arvind Murugan, and Rob Phillips. Proofreading through spatial gradients. Elife, 9:e60415, 2020.

[43] Howard C Berg and Edward M Purcell. Physics of chemoreception. Biophysical journal, 20(2):193– 219, 1977.

[44] Nirmalya Bag, Shuangru Huang, and Thorsten Wohland. Plasma membrane organization of epidermal growth factor receptor in resting and ligand-bound states. Biophysical journal, 109(9):1925–1936, 2015.

[45] Do-Hyeon Kim, Kai Zhou, Dong-Kyun Kim, Soyeon Park, Jungeun Noh, Yonghoon Kwon, Dayea Kim, Nam Woong Song, Jong-Bong Lee, Pann-Ghill Suh, et al. Analysis of interactions between the epidermal growth factor receptor and soluble ligands on the basis of singlemolecule diffusivity in the membrane of living cells. Angewandte Chemie International Edition, 54(24):7028–7032, 2015.

[46] Do-Hyeon Kim, Soyeon Park, Dong-Kyun Kim, Min Gyu Jeong, Jungeun Noh, Yonghoon Kwon, Kai Zhou, Nam Ki Lee, and Sung Ho Ryu. Direct visualization of single-molecule membrane protein interactions in living cells. PLoS Biology, 16(12):e2006660, 2018.

[47] Do-Hyeon Kim, Dong-Kyun Kim, Kai Zhou, Soyeon Park, Yonghoon Kwon, Min Gyu Jeong, Nam Ki Lee, and Sung Ho Ryu. Single particle tracking-based reaction progress kinetic analysis reveals a series of molecular mechanisms of cetuximab-induced EGFR processes in a single living cell. Chemical science, 8(7):4823–4832, 2017.

[48] Marcelo Ehrlich, Werner Boll, Antoine Van Oijen, Ramesh Hariharan, Kartik Chandran, Max L Nibert, and Tomas Kirchhausen. Endocytosis by random initiation and stabilization of clathrin-coated pits. Cell, 118(5):591–605, 2004.

[49] John D Wade, Teresa Domagala, Julie Rothacker, Bruno Catimel, and Edouard Nice. Use of thiazolidine-mediated ligation for site specific biotinylation of mouse EGF for biosensor immobilisation. Letters in Peptide Science, 8:211–220, 2001.

[50] Spandana Kankanala. Binding studies of epidermal growth factor receptor targeted compounds using surface plasmon resonance. PhD thesis, Virginia Commonwealth University, 2009.

[51] AC Myers, JS Kovach, and S Vuk-Pavlović. Binding, internalization, and intracellular processing of protein ligands. derivation of rate constants by computer modeling. Journal of Biological Chemistry, 262(14):6494–6499, 1987.

[52] Boris N Kholodenko, Oleg V Demin, Gisela Moehren, and Jan B Hoek. Quantification of short term signaling by the epidermal growth factor receptor. Journal of Biological Chemistry, 274(42):30169–30181, 1999.

[53] M Zhou, S Felder, M Rubinstein, DR Hurwitz, A Ullrich, I Lax, and J Schlessinger. Real-time measurements of kinetics of EGF binding to soluble EGF receptor monomers and dimers support the dimerization model for receptor activation. Biochemistry, 32(32):8193–8198, 1993.

[54] Teresa Domagala, Nicky Konstantopoulos, Fiona Smyth, Robert N Jorissen, Louis Fabri, Detlef Geleick, Irit Lax, Joseph Schlessinger, William Sawyer, Geoffrey J Howlett, et al. Stoichiometry, kinetic and binding analysis of the interaction between epidermal growth factor (EGF) and the extracellular domain of the EGF receptor. Growth factors, 18(1):11–29, 2000.

[55] Raymond C Harris, Eunkyung Chung, and Robert J Coffey. EGF receptor ligands. The EGF Receptor Family, pages 3–14, 2003.

[56] Jeffrey M Sanders, Matthew E Wampole, Mathew L Thakur, and Eric Wickstrom. Molecular determinants of epidermal growth factor binding: a molecular dynamics study. PloS one, 8(1):e54136, 2013.

[57] Francois Aguet, Costin N Antonescu, Marcel Mettlen, Sandra L Schmid, and Gaudenz Danuser. Advances in analysis of low signal-to-noise images link dynamin and ap2 to the functions of an endocytic checkpoint. Developmental cell, 26(3):279–291, 2013.

[58] Sikao Guo, Alexander J Sodt, and Margaret E Johnson. Large self-assembled clathrin lattices spontaneously disassemble without sufficient adaptor proteins. Biophysical Journal, 121(3):331a, 2022.

[59] Denis S Grebenkov, Ralf Metzler, and Gleb Oshanin. Strong defocusing of molecular reaction times results from an interplay of geometry and reaction control. Communications Chemistry, 1(1):96, 2018.

[60] Ganhui Lan, Pablo Sartori, Silke Neumann, Victor Sourjik, and Yuhai Tu. The energy–speed– accuracy trade-off in sensory adaptation. Nature physics, 8(5):422–428, 2012.

[61] Aidan I Brown and David A Sivak. Theory of nonequilibrium free energy transduction by molecular machines. Chemical reviews, 120(1):434–459, 2019.

[62] Harvey T McMahon and Emmanuel Boucrot. Molecular mechanism and physiological functions of clathrin-mediated endocytosis. Nature reviews Molecular cell biology, 12(8):517–533, 2011.

[63] Daniel Caballero-Díaz, Esther Bertran, Irene Peñuelas-Haro, Joaquim Moreno-Càceres, Andrea Malfettone, Judit López-Luque, Annalisa Addante, Blanca Herrera, Aránzazu Sánchez, Ania Alay, et al. Clathrin switches transforming growth factor-β role to pro-tumorigenic in liver cancer. Journal of Hepatology, 72(1):125–134, 2020.

[64] Guan-Yu Xiao, Aparna Mohanakrishnan, and Sandra L Schmid. Role for ERK1/2-dependent activation of FCHSD2 in cancer cell-selective regulation of clathrin-mediated endocytosis. Proceedings of the National Academy of Sciences, 115(41):E9570–E9579, 2018.

[65] Sidney Redner. A guide to first-passage processes. Cambridge university press, 2001.

[66] Zubenelgenubi C Scott, Aidan I Brown, Saurabh S Mogre, Laura M Westrate, and Elena F Koslover. Diffusive search and trajectories on tubular networks: a propagator approach. The European Physical Journal E, 44(6):80, 2021.

[67] Jenny Ibach, Yvonne Radon, Márton Gelléri, Michael H Sonntag, Luc Brunsveld, Philippe IH Bastiaens, and Peter J Verveer. Single particle tracking reveals that EGFR signaling activity is amplified in clathrin-coated pits. PloS one, 10(11):e0143162, 2015.

[68] Nam Y Lee, Theodore L Hazlett, and John G Koland. Structure and dynamics of the epidermal growth factor receptor C-terminal phosphorylation domain. Protein science, 15(5):1142–1152, 2006.

[69] Theodore R Keppel, Kwabena Sarpong, Elisa M Murray, John Monsey, Jian Zhu, and Ron Bose. Biophysical evidence for intrinsic disorder in the C-terminal tails of the epidermal growth factor receptor (EGFR) and HER3 receptor tyrosine kinases. Journal of Biological Chemistry, 292(2):597–610, 2017.

[70] Li-Zhi Mi, Michael J Grey, Noritaka Nishida, Thomas Walz, Chafen Lu, and Timothy A Springer. Functional and structural stability of the epidermal growth factor receptor in detergent micelles and phospholipid nanodiscs. Biochemistry, 47(39):10314–10323, 2008.

[71] Sarah R Needham, Michael Hirsch, Daniel J Rolfe, David T Clarke, Laura C Zanetti-Domingues, Richard Wareham, and Marisa L Martin-Fernandez. Measuring EGFR separations on cells with ∼10 nm resolution via fluorophore localization imaging with photobleaching. PloS one, 8(5):e62331, 2013.

[72] Nicholas Metropolis and Stanislaw Ulam. The monte carlo method. Journal of the American statistical association, 44(247):335–341, 1949.

